# *BMAL1* Overexpression in Suprachiasmatic Nucleus Protects from Retinal Neurovascular Deficits in Diabetes

**DOI:** 10.1101/2025.02.05.636648

**Authors:** Neha Mahajan, Qianyi Luo, Jodi Lukkes, Surabhi D Abhyankar, Ashay D Bhatwadekar

## Abstract

The suprachiasmatic nucleus (SCN) regulates circadian rhythms and influences physiological and behavioral processes. Disruptions in circadian rhythms (CRD) are observed in type 2 diabetes (T2D), and importantly, CRD acts as an independent risk factor for T2D and its associated complications. BMAL1, a circadian clock gene, is vital for sustaining an optimal circadian rhythm and physiological function. However, the therapeutic potential of BMAL1 overexpression in the SCN to rectify the neurovascular deficits of T2D has yet to be investigated. In this study, db/db mice, a well-established model of T2D exhibiting arrhythmic behavior and the complications of diabetes, were injected stereotaxically with AAV8-Bmal1 or a control virus in the SCN to evaluate the protective effects of correcting the central clock on neurovascular deficits. Given the complex neurovascular network and the eye’s unique accessibility as a transparent system, ocular complications were selected as a model to examine the neuronal functional, behavioral, and vascular benefits of correcting the central clock. BMAL1 overexpression normalized the circadian rhythms, as demonstrated by improvements in the free-running period. The retinal neuronal function improved on electroretinogram, along with optomotor behavior and visual acuity enhancements. Retinal vascular deficits were also significantly reduced. Notably, our approach helped decrease fat content in genetically predisposed obese animals. Since the SCN is known to regulate hepatic glucose production via sympathetic mechanisms, glycemic control, and pyruvate tolerance tests were conducted. Systemically, we observed improved glucose homeostasis in BMAL1-overexpressing mice alongside a substantial reduction in hepatic gluconeogenesis. BMAL1 overexpression lowered plasma norepinephrine and liver TH levels, indicating a protective regulation of adrenergic signaling. Thus, this study underscores the therapeutic potential of targeting circadian clock genes like BMAL1 in the SCN to alleviate metabolic and neurovascular deficits associated with T2D. Our research offers a compelling framework for integrating circadian rhythms into managing diabetes and its complications.

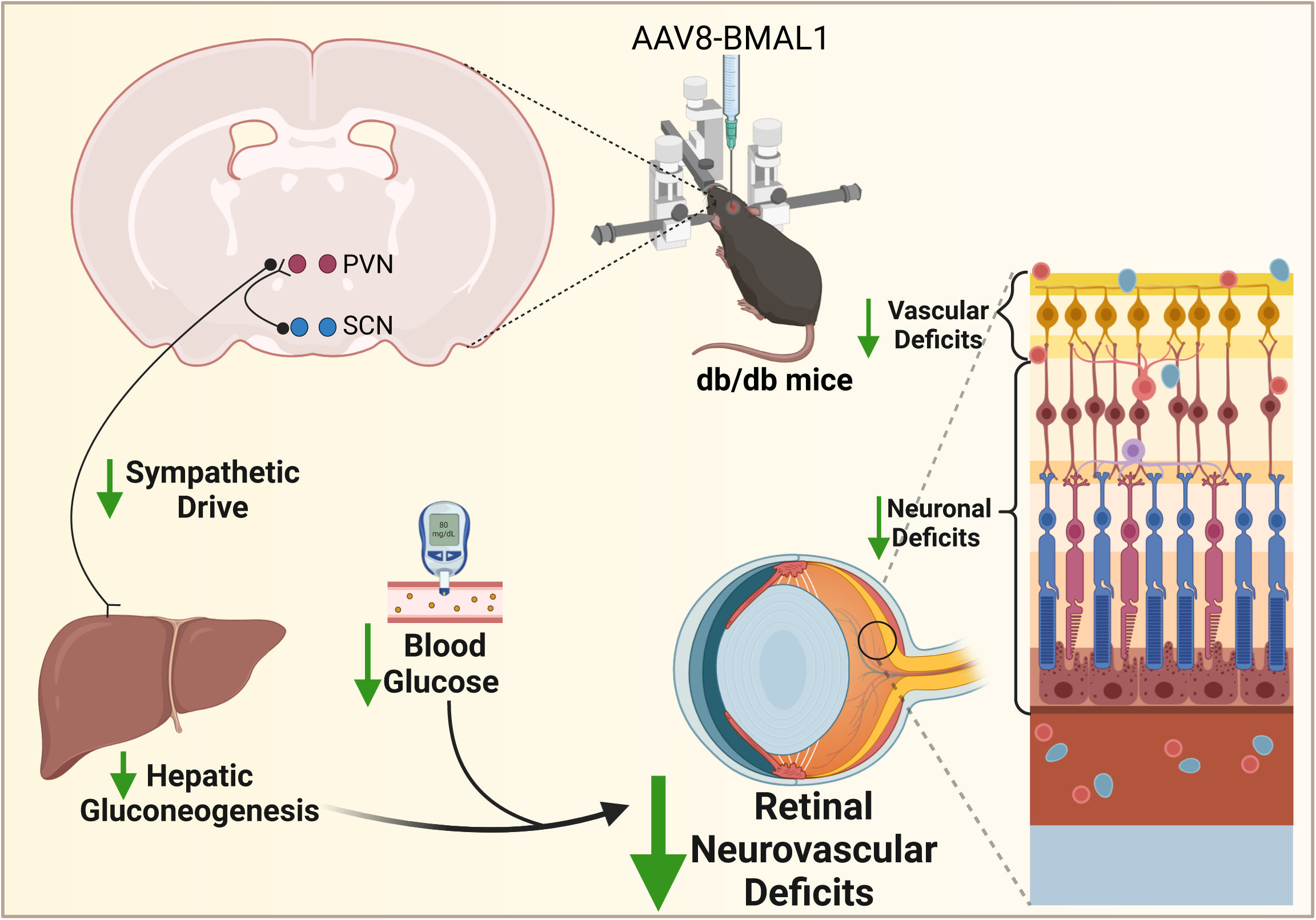

## Introduction

In mammals, a master circadian pacemaker, the suprachiasmatic (SCN) region of the hypothalamus, coordinates the circadian clocks in peripheral tissues to regulate 24-hour rhythms of physiological functions [1]. Transcriptional and translational feedback loops drive these circadian oscillations at the cellular level. The transcription factor brain and muscle ARNT-like protein 1 (*BMAL1*) interacts with circadian locomotor output cycles kaput (*CLOCK*) gene to form a heterodimer, eventually to mediate the transcription of downstream negative regulators Period (*Per*) and Cryptochrome (*Cry*) genes. The *Per/Cry* heterodimer inhibits the expression and transcriptional activity of *BMAL1*/*CLOCK* genes [2]. The fine-tuning and strict regulation of this circadian feedback loop is critical for circadian rhythmicity in mammals.

*BMAL1* is the only non-redundant gene in the central circadian clock, elevating its significance in various circadian rhythm-focused studies [3]. Interestingly, several studies have linked the *BMAL1* abnormal expression patterns to metabolic complications, including type 2 diabetes (T2D). *BMAL1* deletion has been shown to promote T2D, obesity, lipogenesis, and pancreatic β cell impairment [4–6], and conversely, pancreatic β-cell-specific overexpression of *BMAL1* was found to be protective against obesity-induced glucose intolerance, and mice-overexpressing *BMAL1* in the same study had better glucose-stimulated insulin release [7]. This growing evidence reflects the significant potential of *BMAL1* in chronic diabetes.

With the increasing prevalence of diabetes worldwide, the associated microvascular complications are growing rapidly among young, working-age adults [8, 9]. The eye is uniquely positioned to study neurovascular complications of diabetes due to the combination of a transparent window and an intricate network of retinal neurons and vasculature. Additionally, it exhibits the most common complication of diabetes, diabetic retinopathy. The mammalian retina also has an autonomous circadian clock system independent of the SCN [10]; previously, we reported that circadian clock disruption negatively affects visual function in mice [11]. Several studies have shown the adverse effects of BMAL1 deletion on retinal cone and rod cells, which could accelerate retinal microvascular and macrovascular injuries [12–14]. These findings strengthen our premise for using the eye as a model system to assess the therapeutic benefits of correcting the circadian clock. We employ a novel therapeutic strategy for correcting overall circadian rhythm by overexpressing clock gene BMAL1 centrally in the SCN via stereotaxic delivery. We provide evidence that *BMAL1* overexpression improves neuronal function and reduces acellular capillaries, i.e., vascular deficits of diabetes, by correcting systemic glucose metabolism. Importantly, the beneficial effects of *BMAL1* overexpression were associated with decreased hepatic gluconeogenesis and noradrenaline signaling pathways. These observations implicate the potential therapeutic targeting of circadian clock genes for diabetes and its complications.

## Research Design and Methods

### Animals

The B6.BKS(D)-Lepr^db^/J (an animal model for type 2 diabetes; db/db) and Lepr^db^/^+^ db/m (heterozygotes; db/m) mice [stock number 000697] were procured from the Jackson Laboratory (Bar Harbor, ME, USA) and housed in the animal care facility at Glick eye institute, Indiana University. All the animals were kept under normal physiological conditions (12-hour light/dark conditions), with free access to food and water *ad libitum.* All the experiments performed were per the Guiding Principles in the Care and Use of Animals (National Institutes of Health) and the Association for Research in Vision and Ophthalmology’s Statement for the Use of Animals in Ophthalmic and Vision Research.

### AAV-*BMAL1* delivery to the suprachiasmatic nucleus (SCN)

AAV-*BMAL1* was designed and procured from the Ocular Gene Therapy Core at the University of Florida, Gainesville, FL. The *BMAL1* was cloned with smCBA promoter for ubiquitous expression with a GFP tag and subsequently packaged in AAV-8 for delivery and expression as smCBA-mBmal1-P2A-GFP, the AAV8-smCBA-GFP was used as a control virus. We performed a stereotaxic delivery of the AAV-*BMAL1* vector using coordinates of bregma -0.96, lateral 0.5 mm, and ventral 9 mm for SCN delivery in 8-week-old mice. To validate the SCN deliveries, three mice were sacrificed four weeks post-stereotaxic surgeries followed by sectioning and staining. After successfully validating the SCN deliveries (**Suppl Fig 1**), the mice were divided into the following groups: 1) db/m + NV (control mice/ No virus), 2) db/m + Cont (db/m with AAV only), 3) db/m + *BMAL* (db/m with AAV-*BMAL1*), db/db + NV (diabetic control mice/ No virus), 2) db/db + Cont (db/db with AAV only), 3) db/db + *BMAL* (db/db with AAV-*BMAL1*). All the mice were maintained for six months post-injections for the following listed experiments.

### Echo MRI

Live unanaesthetised mice are guided into a plastic tube designed to restrict their movement but not constrain them. The animals are then subjected to a 1-4 min imaging period in the gantry using a low energy (0.05T) electromagnetic field. Based on emitted T1 and T2 relaxation curves, lean mass, fat mass, free water, and total body weight are calculated using standard algorithms.

### Wheel Running Activity

Wheel running activity was measured using our previously reported method [11]. All the animals were housed individually in running wheel cages in sound attenuated and ventilated isolation cabinets (Phenome Technologies, Chicago, IL, USA) for 14 weeks. Wheel running activity was recorded every minute for 10 weeks using Actimetrics (Actimetrics, Chicago, IL, USA) hardware and monitored and analyzed using Clocklab (Actimetrics, Chicago, IL, USA). The mice were exposed to 12H light and 12H dark for the first 14 days and subsequently kept in the constant dark to assess the free-running activity rhythms. The actogram data from 15 to 70 days were used to determine the period and quantify wheel running activity using Clocklab. The period was determined by periodogram analysis (L12:D12). Wheel running activity counts were determined by averaging the total activity during light and dark phases and were expressed for 24 hours to make them comparable between groups.

### Neuronal Funciton using Electroretinogram (ERG)

The retinal function was measured in the animals employing dark-adapted/scotopic ERG (LKC Technologies, Inc, Gaithersburg, MD, USA). All the animals were dark-adapted for 24 hours before proceeding with the ERG recordings. The mice were anesthetized by administering an i.p. injection of ketamine (100 mg/kg) and xylazine (5 mg/kg). The pupils were dilated using a topical application of 1% tropicamide and 2.5% phenylephrine (Alcon laboratories). The eyes were kept moist using a 2.5% hypromellose ophthalmic demulcent solution/Gonak (Akorn). For the ERG recordings, the ground needle electrode was placed on the base of the tail, and the reference electrode was sub-dermally placed between the eyes. The gold loop electrodes (LKC Technologies, Inc, Gaithersburg, MD, USA) placed over the cornea were used to record ERG response. The stimulus flash intensities of 0.025, 0.25, and 2.5 cd.s/m^2^ for scotopic conditions were presented in a UTAS ganzfeld illuminator (LKC Technologies). The values for a wave and b wave amplitudes and their implicit times were obtained from an inbuilt analysis tool by LKC Technologies.

### Optomotor Response Behavior (OMR)

The spatial vision was quantified in the animals by detecting the spatial frequency threshold of optometer response behavior using an OptoMotry device (CerebralMechanics, Inc.). Tracking head movements in response to rotating sine wave gratings (100% contrast) were recorded in free moving mice. Spatial frequency was systematically increased in a staircase method until the animal did not respond, and the highest spatial frequency the animal could track was identified as the threshold. The threshold obtained for each eye was reported.

### Immunohistochemistry for Tyrosine Hydroxylase

The liver samples were fixed using 4% PFA in PBS for 24 hours, followed by paraffin embedding. The sections were placed on charged slides and dried at 56°C overnight. Slides were subsequently deparaffinized in xylene and hydrated through descending grades of ethyl alcohol to distilled water, and placed in Tris Buffered Saline pH 7.4 (Scytek Labs, Logan, UT) for 5 minutes for pH adjustment. The sections were then subjected to enzyme-induced epitope retrieval in 0.03% Pronase E/TBS (Millipore Sigma/Scytek) in an incubator at 37°C for 10 minutes, followed by several rinses in distilled water. Before proceeding with blocking for non-specific proteins with rodent Block M (Biocare, Concord, CA) for 20 minutes, the sections were pre-treated with 3% hydrogen peroxide/methanol for 30 minutes at 25°C, rinsed with distilled water, and washed with TBST for 5 minutes, followed by micro-polymer staining performed at room temperature on the Biocare intelliPATH automated stainer. Lastly, the sections were incubated with primary antibody (rabbit Tyrosine Hydroxylase (Millipore Sigma, Temecula, CA)) at 1:150 in Normal Antibody Diluent (Scytek) for 1 hour, followed by rodent HRP Polymer (Biocare) incubated for 30 minutes. Reaction development utilized Romulin AEC (Biocare) for 5 minutes, counterstained in CATHE Hematoxylin diluted 1:10 for 1 minute, followed by air drying, dipping in xylene, and coverslipping with permanent mounting media. The slides were imaged at 20X under a fluorescent microscope (Zeiss AXIO Observer.A1 Inverted Fluorescence Microscope, Carl Zeiss MicroImaging GmbH).

### Vascular deficits

Animals were euthanized, and the eyes were enucleated and fixed in 4% paraformaldehyde. A day before trypsin digestion, the retinas were isolated following the previously reported procedure [15]. The isolated retina was placed in 50 mL water for unfixing overnight. The individual retina was incubated in 3% trypsin at 37°C for 2 hours the next day. The trypsin-digested retina was placed in a Petri dish, and the internal limiting membrane was gently separated from the peripheral retina with fine forceps. Then, using Vannas scissors, the internal limiting membrane was isolated from an optic nerve. Subsequently, the neural retina was removed, and the isolated retinal vasculature was stained with periodic acid and Schiff’s base to assess acellular capillary numbers.

### Norepinephrine levels using ELISA

Whole blood was collected in an EDTA-coated collection tube before sacrificing the animals. After 30 minutes, the blood samples were centrifuged at 2000 x g for 20 minutes. The clear supernatant or plasma was separated and stored at -80° C. Plasma norepinephrine levels were quantified using a commercially available norepinephrine ELISA kit [cat no. 3836; Novus Biologicals LLC, CO, USA] as per the manufacturer’s guidelines.

### Glucose Tolerance Test (GTT)

The animals were fasted for 4 hours, and basal blood glucose levels were monitored using a commercially available glucometer (AlphaTrek2) to set up basal or zero-time blood glucose. After which, glucose solution (1g/kg/bw) was administered *via* the i.p. route. The blood glucose was then measured again at 10, 20, 30-, 60-, 90-, and 120 minutes post-glucose administration.

### Insulin Tolerance Test (ITT)

The animals were fasted for 2 hours for ITT, and basal blood glucose levels were quantified as described above. An i.p. injection of 0.5 IU/kg insulin Humulin R U-100 was administered to the animals, followed by measurement of glucose levels as described in GTT.

### Pyruvate Tolerance Test (PTT)

The animals were fasted for 16 hours before basal blood glucose measurements and sodium pyruvate (P5280 Millipore Sigma, US) prepared in sterile PBS @ 1g/kg/bw was given intra-peritoneally. The blood glucose levels were afterward quantified using a glucometer for 15, 30, 60, 90, and 120 minutes as described earlier.

### Statistical Significance

All the data were expressed as Mean ± SEM. The data was analyzed on GraphPad Prism V.10.0.0 for Windows (San Diego, California; www.graphpad.com) using either one-way ANOVA or Brown Forsythe and Welch’s ANOVA test. Data were considered statistically significant when the p-value was less than 0.05.

## Results

### 1. *BMAL1* overexpression improved free-running periods in db/db mice

After *BMAL1* overexpression in the SCN of db/db mice, we first investigated the voluntary free wheel running activity (**Fig 1A)**. We observed that db/db mice had reduced total activity compared to the db/m mice **(suppl Fig 2)**, and there was no effect of *BMAL1* overexpression. The free running periods were significantly delayed in the db/db mice and db/db mice with the control virus **(Fig 1B)**. Interestingly, db/db mice overexpressing *BMAL1* had significantly improved free-running period **(Fig 1B)**, suggesting that *BMAL1* overexpression corrects the endogenous circadian rhythms in db/db mice.

**Fig 1:**
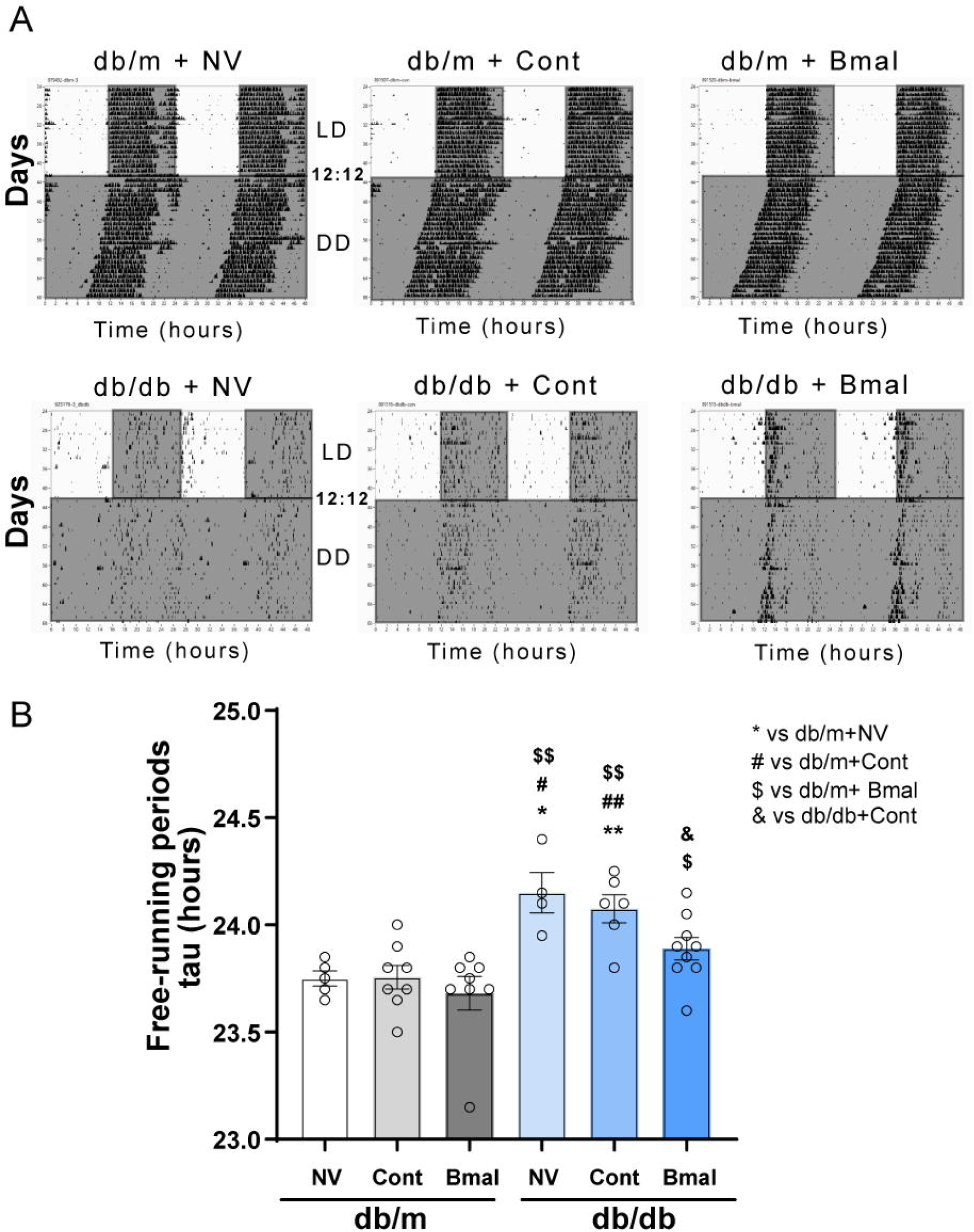
SCN-*BMAL1* overexpression improves free-running periods. **(A)** Representative double-plotted actograms of wheel running activity under 12 hr. light and dark (LD12:12) and constant dark (DD) conditions. **(B)** Bar graphs presenting the free-running period of the respective groups. N: db/m+NV-5; db/m+cont-8; db/m+*BMAL* -8; db/db+NV-4; db/db+Cont-6; db/db+ *BMAL* -9. The data is presented as Mean ± SEM and analyzed using Brown Forsythe and Welch’s ANOVA test where different symbols signify the following: * vs db/m + NV, *=p<0.05, ** = p<0.01; # vs db/m + Cont, # p<0.05, ## p<0.01; $ vs db/m + *BMAL*, $ p<0.05, $$ p<0.01 and & vs db/db + Cont, p<0.05.

### 2. *BMAL1* overexpression improved OMR in db/db mice

As we observed an improvement in the endogenous circadian rhythms of db/db mice, we studied the beneficial effects of central *BMAL1* overexpression optomotor behavior. Firstly, we studied OMR tracking to check whether *BMAL1* influences visual performance. The spatial frequency threshold was significantly reduced in the db/db and db/db + Cont, which was significantly improved in the db/db mice overexpressing *BMAL1* **(Fig 2A)**. The contrast sensitivity was significantly lower in the db/db mice irrespective of *BMAL1* overexpression compared to the db/m groups **(Fig 2B).**

**Fig 2:**
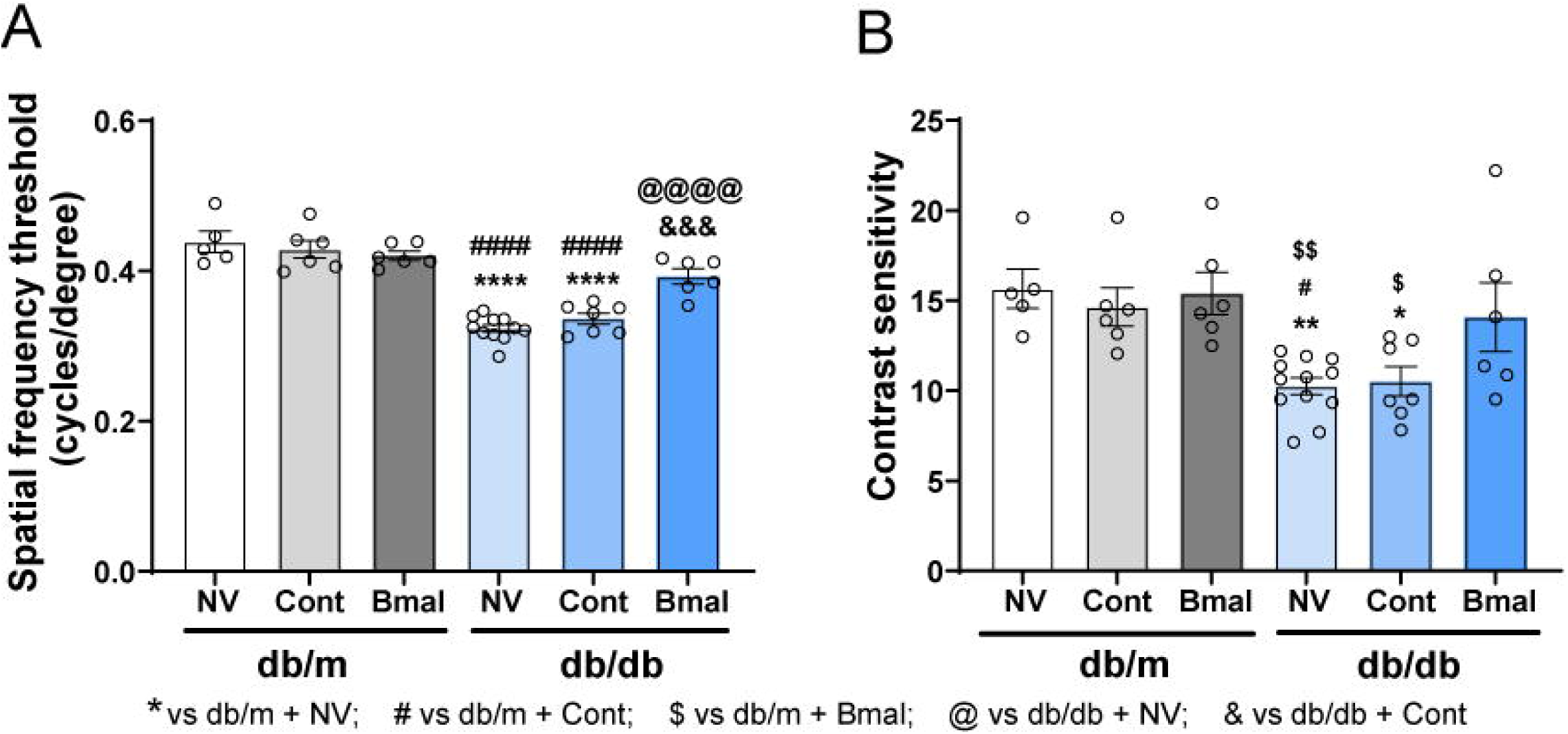
SCN-*BMAL1* overexpression improved visual acuity behavior. **(A)** db/db mice overexpressing Bmal1 showed a significant increase in the spatial frequency threshold measured using an optomotor reflex tracking System. **(B)** Contrast sensitivity remained unchanged with the overexpression of *BMAL1*. N= db/m+NV-5; db/m+cont-6; db/m+ *BMAL* -6; db/db+NV-12; db/db+Cont-7; db/db+ *BMAL* -6. The data is presented as Mean ± SEM and analyzed using One-way ANOVA followed by Tukey’s post-hoc test, * vs db/m + NV, * = p<0.05, ** = p<0.01, *** = p<0.001; # vs db/m + Cont, # = P<0.05, ### = p<0.001; $ vs db/m + *BMAL*, $ = p<0.05, $$ = p<0.01; @@@ vs db/db + NV, p<0.001 and &&& vs db/db + Cont, p<0.001.

### 3. *BMAL1* overexpression improved retinal function in db/db mice

Further, we examined the effect of *BMAL1* overexpression on retinal function using an electroretinogram (ERG). Mice were subjected to scotopic ERG to evaluate bipolar cells and photoreceptor cells’ activity. There was a significant reduction in the b-wave amplitude in the db/db and db/db + Cont group, which was improved under *BMAL1* overexpression in db/db mice, suggesting a protective effect of *BMAL1* overexpression on bipolar cells in db/db mice **(Fig 3A).** Similarly, there was also a reduction in the b-wave peak latency time under *BMAL1* overexpressing db/db mice **(Fig 3B)**. The a-wave amplitude and peak latency time, reflecting rod cells’ activity, were also improved in db/db mice overexpressing *BMAL1*, but the difference was significant only at the 2.5 log cd.s/m^2^ flash intensity **(Fig 3C&D)**. The ERG analysis suggested that *BMAL1* overexpression helps improve retinal function *by improving bipolar cells and rod photoreceptors’* functions.

**Fig 3:**
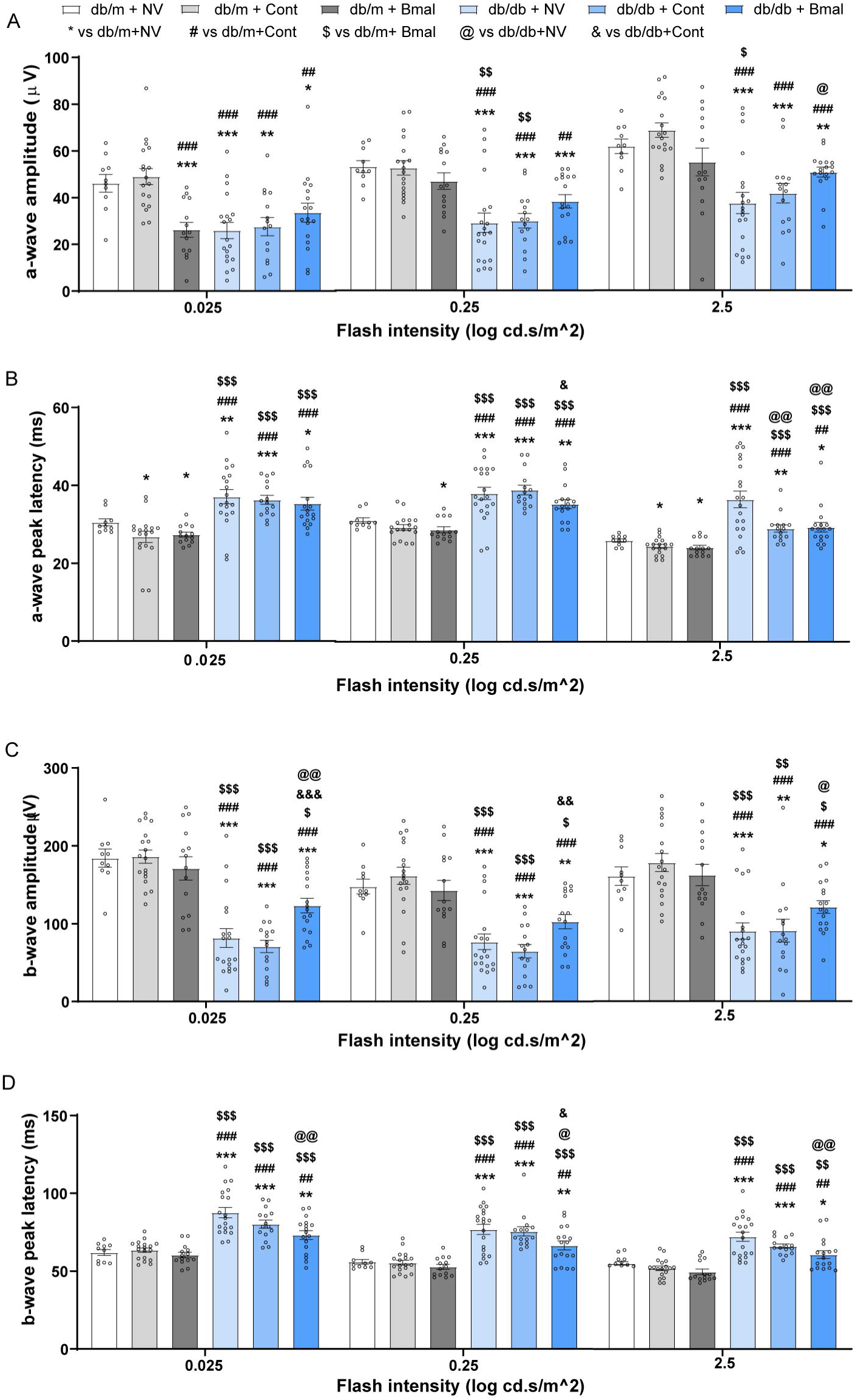
SCN-*BMAL1* overexpression improved neuronal function. **(A)** Scotopic a-wave quantification (**B**) a-wave peak latency time **(C)** Scotopic b-wave quantification **(D)** b-wave implicit time quantification. N: db/m+NV-10; db/m+cont-18; db/m+ *BMAL* -14; db/db+NV-19; db/db+Cont-15; db/db+ *BMAL* -17. The data is presented as Mean ± SEM and analyzed using Brown Forsythe and Welch’s ANOVA test; *p<0.05, **p<0.01, **p<0.001.

### 4. *BMAL1* overexpression improved the vascular deficits in db/db mice

After examining the behavioral functional vision using OMR and retinal functions using ERG, we next examined the vascular phenotype (**Fig 4**). The db/db mice and db/db + Cont showed a significant increase in the acellular capillaries compared to the db/m groups (**Fig 4A and 4B**). The *BMAL1* overexpression showed a significant reduction in the number of acellular capillaries in db/db mice, suggesting a preventive effect on DR (**Fig 4B**).

**Fig 4:**
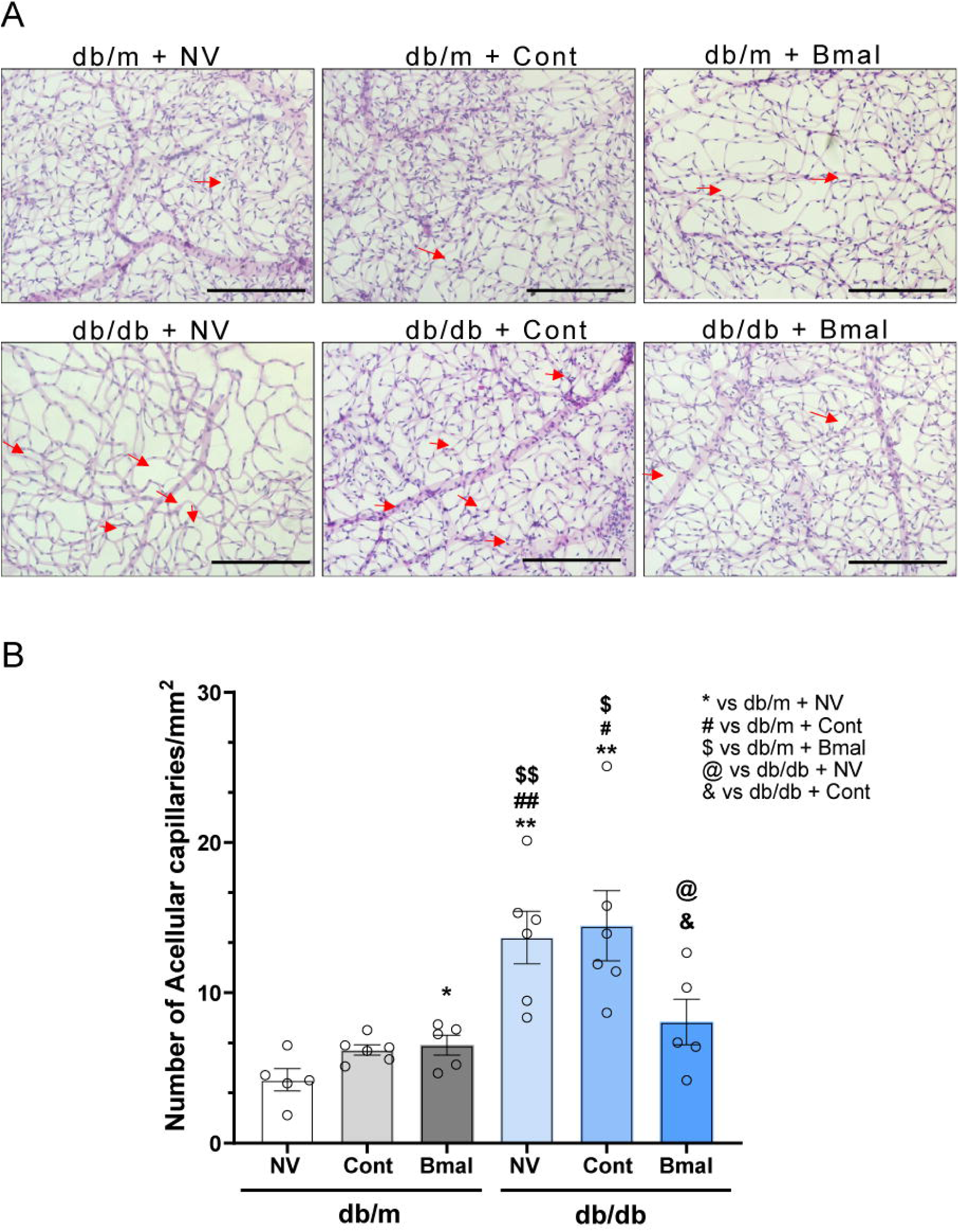
A decrease in the vascular deficits in SCN-*BMAL1* overexpressed db/db mice. **(A)** Representative images of trypsin-digested retinas from the respective groups and red arrows showing the changes in acellular capillary numbers. **(B)** Bar chart showing the quantification for the same. N= 5, Magnification 20X and Scale Bar 100µM; The data is presented as Mean ± SEM and analyzed using Brown-Forsythe and Welch ANOVA test; * vs db/m + NV, *p<0.05, **p<0.01; # vs db/m + Cont, #p<0.05, ##p<0.01; $ vs db/m + *BMAL*, $ p<0.05, $$ p<0.01; @ vs db/db + NV, p<0.05 and & vs db/db + Cont, p<0.05.

### 5. *BMAL1* overexpression improved glucose homeostasis and physiological parameters in db/db mice

While circadian rhythms, neurovascular deficits, and behavioral responses were corrected using our therapeutic strategy of correcting the central clock, the systemic effects of *BMAL1* overexpression were not assessed. Reported literature suggests that *BMAL1* knock-out animals exhibit weight gain and glucose intolerance [16, 17]. To investigate the peripheral beneficial outcomes of central *BMAL1* overexpression, we first evaluated the body composition of mice using EchoMRI **(Fig 5A-C)**. The body weight was significantly higher in db/db mice, while the *BMAL1* overexpression led to a decrease in body weight when compared to the dbdb-control group **(Fig 5A),** the difference remained statistically insignificant. Strikingly, the fat mass was reduced significantly in *BMAL1* overexpressing db/db mice compared to db/db mice. There was no observable difference in lean mass percentage among all the groups **(Fig 5C)**.

**Fig 5:**
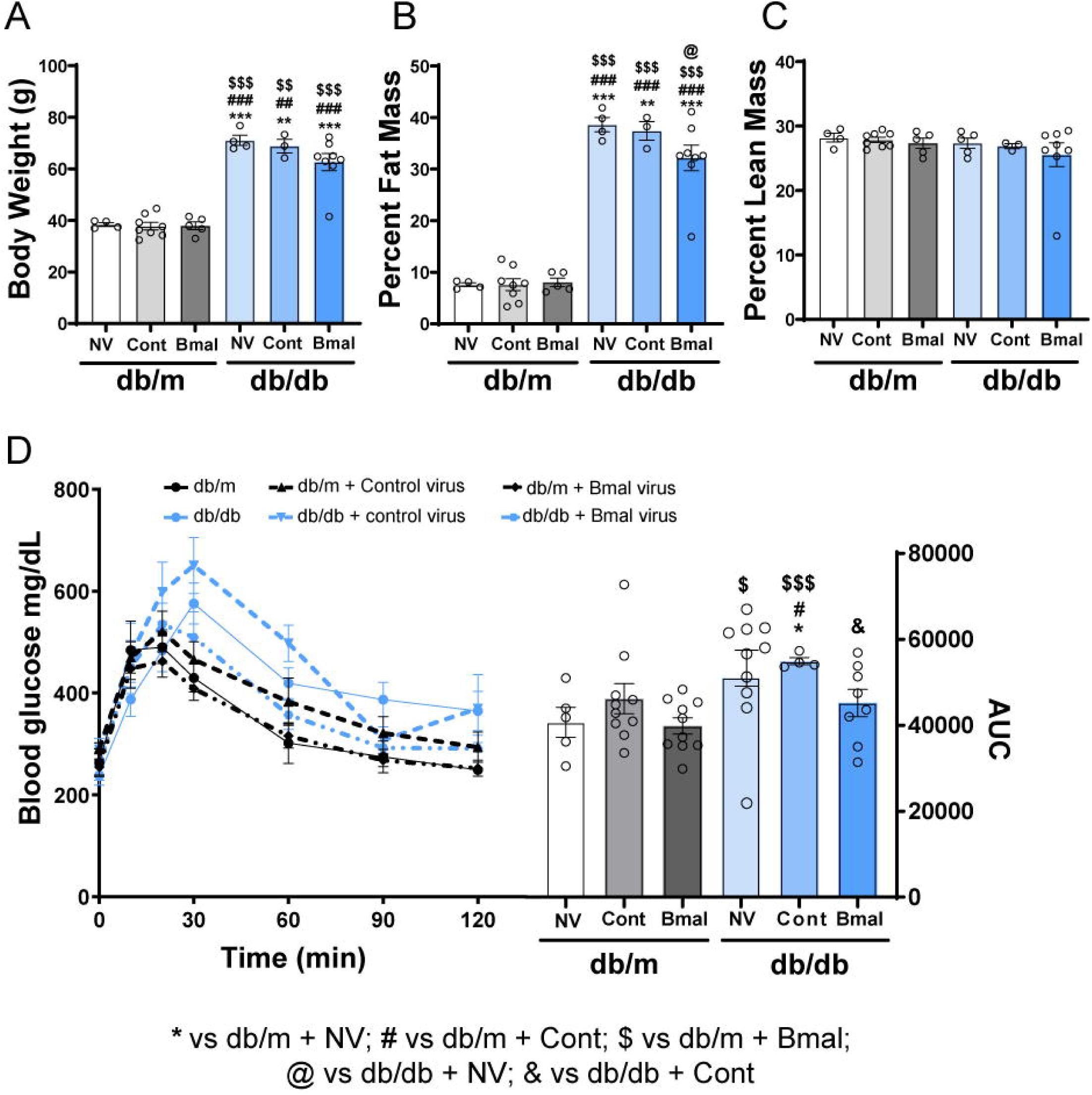
Effect of SCN-*BMAL1* on anthropometry and glucose homeostasis. (**A**) Body weight (**B**) Fat mass and (**C**) Lean mass measured using Echo MRI. (**D**) GTT and its AUC in respective groups. N for A-C: db/m+NV-4; db/m+cont-8; db/m+ *BMAL* -5; db/db+NV-4; db/db+Cont-3; db/db+ *BMAL* -8. N for D: db/m+NV-5; db/m+cont-10; db/m+ *BMAL* -10; db/db+NV-12; db/db+Cont-4; db/db+ *BMAL* -8. The data is presented as Mean ± SEM and analyzed using Brown-Forsythe and Welch ANOVA test; * vs. db/m + NV, *p<0.05, **p<0.01, ***p<0.001; # vs. db/m + Cont, #p<0.05, ##p<0.01, ###p<0.001; $ vs db/m + *BMAL*, $ p<0.05, $$ p<0.01, $$$ p<0.001; @ vs db/db + NV, p<0.05 and & vs db/db + Cont, p<0.05.

Next, with the help of GTT, the effect of *BMAL1* overexpression on glucose homeostasis was evaluated. A higher AUC in db/db and db/db + Cont mice reflected an impaired glucose clearance, which was improved after *BMAL1* overexpression in db/db mice **(Fig 5D)**. We also examined insulin sensitivity using ITT, which showed a reduction in blood glucose clearance in db/db mice, and *BMAL1* overexpression was unable to improve insulin-dependent glucose homeostasis in db/db mice **(Suppl Fig 3).**

### 6. *BMAL1* overexpression improved hepatic gluconeogenesis through norepinephrine signaling

*BMAL1* is a well-known regulator of the central circadian clock. From our GTT, we observed an improvement in glucose intolerance. However, the insulin-dependent glucose homeostasis remains unchanged. Alternatively, *BMAL1* also regulates hepatic glucose metabolism [18]. To ascertain whether similar mechanisms play a role in our study’s beneficial effects of central *BMAL1* overexpression, we first performed a PTT, a routinely used test for hepatic gluconeogenesis. We found an increase in hepatic glucose production/gluconeogenesis in db/db, and db/db + Cont mice, which was significantly reduced under *BMAL1* overexpression **(Fig 6A)**, suggesting a direct impact of SCN *BMAL1* on hepatic gluconeogenesis and glucose metabolism.

**Fig 6:**
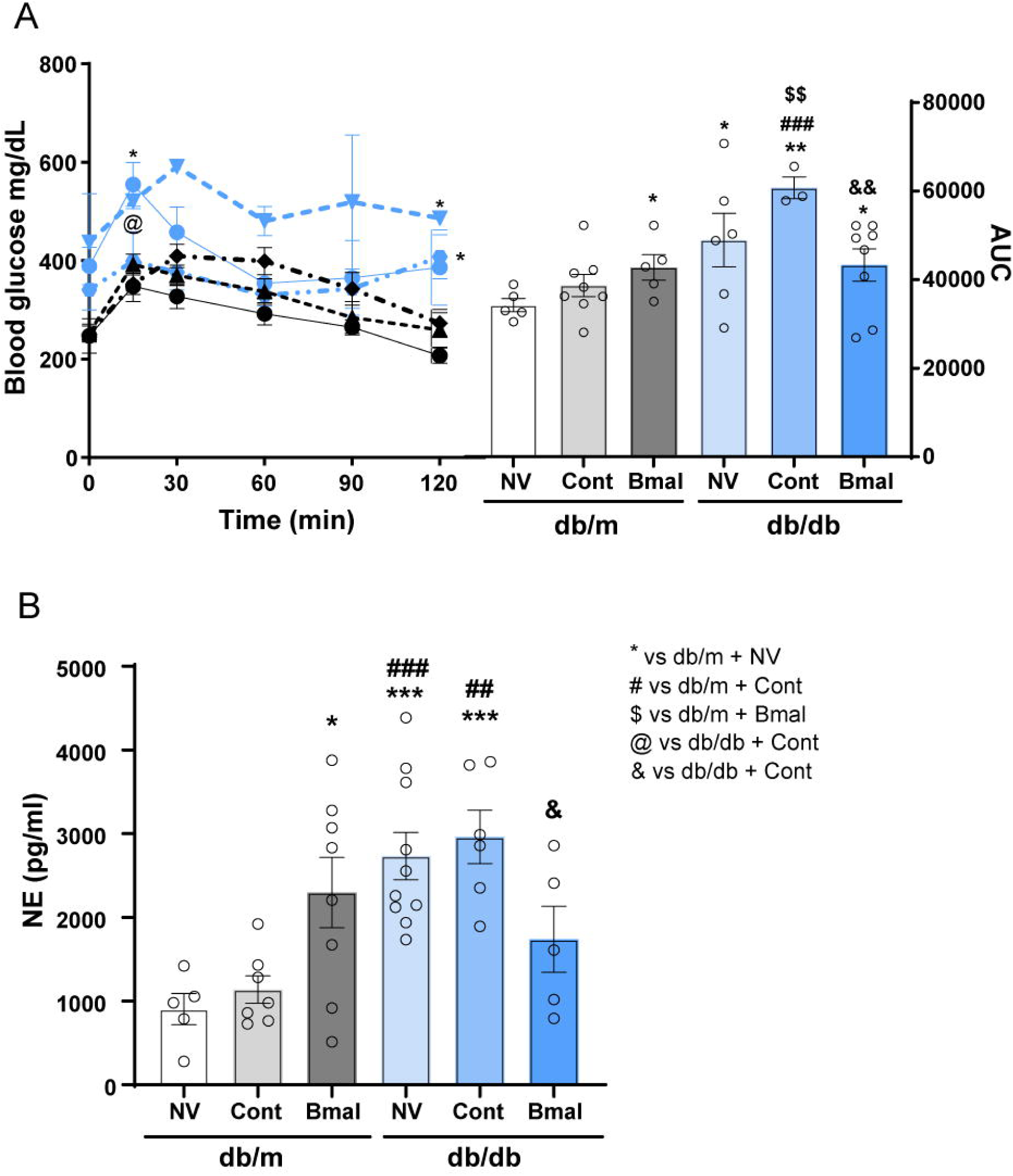
Effect of SCN-*BMAL1* overexpression on hepatic gluconeogenesis and sympathetic nervous system. (**A**) Pyruvate tolerance test (PTT) and its AUC in respective groups. (**B**) Plasma norepinephrine (NE) levels were quantified using ELISA. N for PTT: db/m+NV-5; db/m+cont-8; db/m+*BMAL*-5; db/db+NV-6; db/db+Cont-2; db/db+*BMAL*-8. N for NE measurement: db/m+NV-5; db/m+cont-7; db/m+ *BMAL* -8; db/db+NV-10; db/db+Cont-6; db/db+ *BMAL* -5. The data is presented as Mean ± SEM and analyzed using Brown Forsythe and Welsch’s ANOVA test; * vs db/m + NV, *p<0.05, **p<0.01,**p<0.001; # vs db/m + Cont, ##p<0.01, ###p<0.001; $$ vs db/m + *BMAL1*, p<0.01; and & vs db/db + Cont, & p<0.05, && p<0.01.

Interestingly, hyperglycemia and impaired glucose homeostasis can elevate the sympathetic drive, increasing the norepinephrine levels [19]. Reciprocally, literature also suggests that poorly controlled diabetes could lead to an increase in plasma norepinephrine levels [20, 21]. We were intrigued to investigate if central overexpression of *BMAL1* could work through the sympathetic drive. So, we quantified plasma norepinephrine levels to understand the mechanism behind *BMAL1* peripheral protective effects on glucose homeostasis. The norepinephrine levels were significantly higher in db/db mice, and *BMAL1* overexpression in db/db mice reduced the circulating norepinephrine levels **(Fig 6B).**

Moreover, we found an upregulation for tyrosine hydroxylase (TH) in the liver sections of db/db and db/db+Cont groups **(Fig 7)**. TH is a rate-limiting enzyme in the synthesis of norepinephrine, therefore affecting the adrenergic pathways [22, 23]. The immunohistochemistry experiment revealed that db/db mice activated the sympathetic nervous system, which was significantly regulated in *BMAL1* overexpressing db/db mice (**Fig 7**).

**Figure 7:**
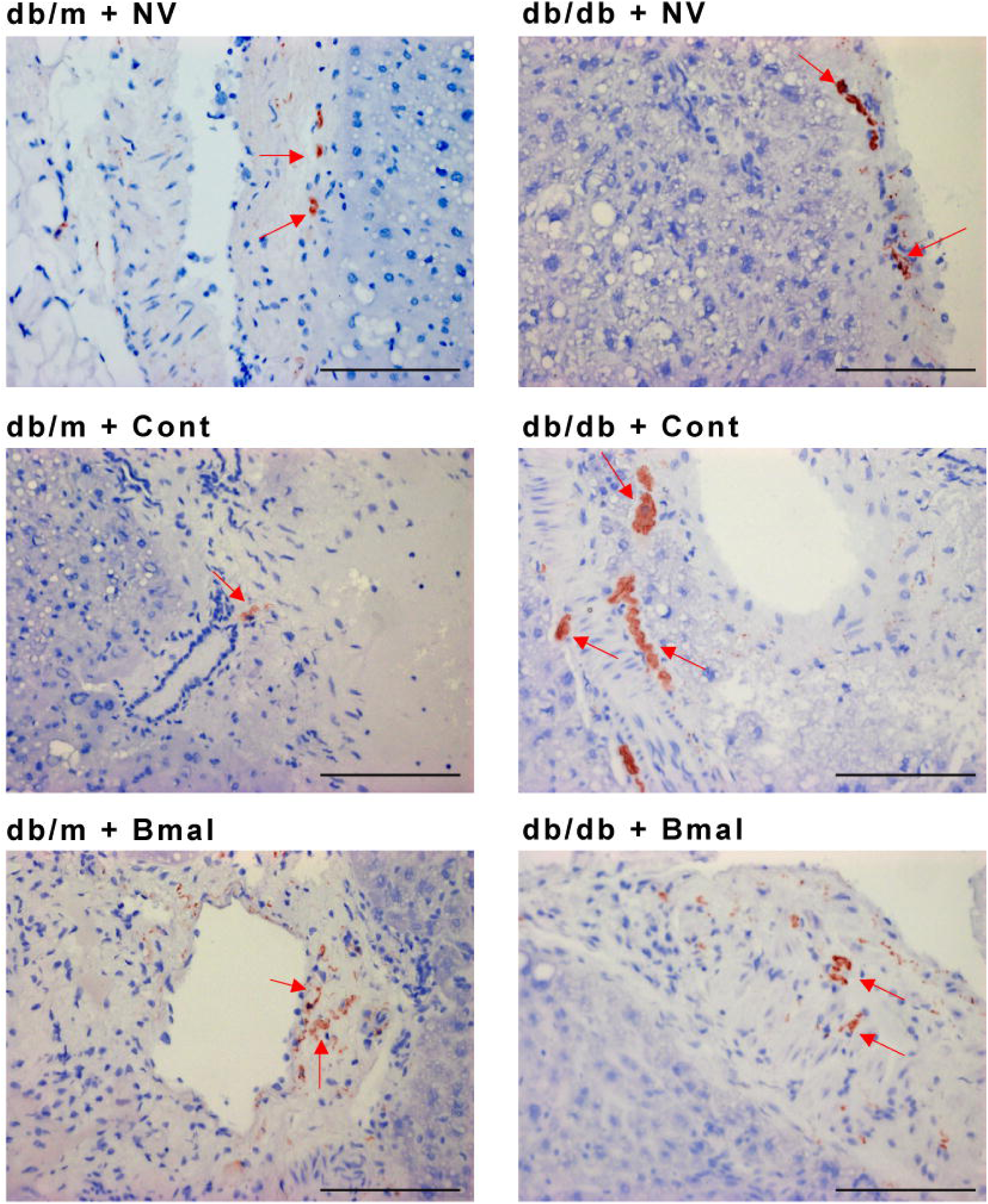
SCN-*BMAL1* overexpression downregulates Tyrosine hydroxylase (TH) expression. Representative images of TH-stained liver sections. Magnification: 20X and Scale Bar 100µM. N= 5

In conclusion, our data suggested that the sympathetic nervous system was activated in db/db mice, resulting in hepatic gluconeogenesis stimulation, which led to dysregulation of glucose homeostasis. *BMAL1* overexpression in db/db mice prevented all these complications.

## Discussion

Disruptions in circadian rhythms are linked to various metabolic disorders like diabetes, yet there has been little exploration of therapeutic strategies aimed at the circadian clock. This study highlights the protective role of BMAL1 gene overexpression in the SCN, impacting circadian rhythms, neurovascular deficits, and systemic physiological aspects in T2D. Furthermore, our findings offer distinctive mechanistic insights, illustrating how BMAL1 overexpression in the SCN influences sympathetic activity and glucose intolerance, a relationship that hasn’t been previously investigated.

In mammals, the circadian rhythms are regulated by a master circadian clock located in the SCN of the hypothalamus, which in turn governs the peripheral circadian clock of peripheral organs. Interestingly, recent research has shown the downregulation of hippocampal *BMAL1* in streptozotocin/high-fat diet-induced diabetic mice [24]. To facilitate the improvement in the central circadian clock, we injected AAV-*BMAL1* in the SCN of db/db mice. The overexpression of *BMAL1* in db/db mice improved the free-running period. This observation is consistent with the previous studies where *BMAL1* deletion in arginine vasopressin neurons (AVP) of the dorsal SCN region lengthened the free-running period in AVP-*BMAL1* knock-out mice [25].

Research has shown that diabetic mice experience a reduction in the expression of clock genes [26–28], which contributes to circadian arrhythmicity observed in db/db mice and leads to disrupted glycemic control [11, 29, 30]. Our findings indicate that by overexpressing *BMAL1* in the SCN of diabetic mice, we could correct the central circadian clock. This intervention halted the rise in acellular capillaries and vascular deficits, which are typically hallmarks of the complications of diabetic retinopathy. Also, the reduction in b-wave amplitude in db/db mice was improved after *BMAL1* overexpression in the SCN of db/db mice. Indeed, conditional *BMAL1* knock-out mice also display retinal deficits, explicitly affecting the circadian rhythmicity of ERG b-wave amplitude [31]. However, in our study, we do not expect *BMAL1* overexpression in SCN to directly impact retinal *BMAL1* and influence ocular parameters, though we cannot exclude the potential for improved retrograde signaling. This intriguing possibility warrants further investigation in future studies.

One of our study’s notable findings is a decrease in hepatic glucose production as the mode of action, which was not studied earlier in this context. The liver is a principal organ for glucose storage, and disruption in liver functions has detrimental metabolic consequences. Previously, injecting the transneuronal pseudorabies virus into the liver resulted in retrograde labeling of CNS neurons. Notably, the localization of third-order neurons in the SCN illustrates anatomical pathways that enable the biological clock to influence autonomic input to the liver, emphasizing the direct effects of restoring the SCN clock on liver function [32]. Moreover, sympathetic activation stimulates hepatic glucose production through gluconeogenesis and glycogenolysis, contrary to the effects of parasympathetic activation, leading to a reduction in glucose production [33, 34]. Intriguingly; we observed an increase in the plasma levels of norepinephrine, a sympathetic neurotransmitter in db/db mice, and *BMAL1* overexpressing db/db mice had a significant reduction in NE levels, suggesting impaired gluconeogenesis and glucose homeostasis in db/db mice.

Moreover, as we delve deeper into the sympathetic system and hepatic gluconeogenesis as a principal model of action for the SCN’s effects, we also analyze the hepatic expression of the tyrosine hydroxylase (TH) enzyme. TH is a rate-limiting enzyme for synthesizing catecholamines (epinephrine, norepinephrine, and dopamine), known to be expressed in the nerve fibers around the portal vein, bile duct, or hepatic arteries of the liver. Consistent with the reported literature for a higher expression of TH enzyme in the liver sections of obese animals [35], we found a similar trend in db/db mice. However, with the overexpression of *BMAL1* in db/db mice, the expression of TH was significantly decreased. We speculate that a higher hepatic TH expression with a simultaneous increase in the plasma norepinephrine levels leads to impaired glucose metabolism in diabetic mice. The correction of the central circadian clock *via* overexpressing SCN-*BMAL1* has profound effects against peripheral impaired glucose homeostasis. So far, published literature has shown that NE can influence rodent livers’ circadian rhythms and clock gene expression [36].

Although the central overexpression of BMAL1 enhanced glycemic control, it did not impact insulin-dependent glucose clearance. This lack of change may be due to the genetic background of db/db mice (which are deficient in leptin receptors), as BMAL1 may interact with leptin and its receptors to influence various metabolic pathways, including insulin sensitivity and weight gain. [37]. It’s also important to emphasize that overexpressing BMAL1 in the SCN reduced fat mass in genetically obese mice. With recent advancements in anti-diabetic treatments (such as GLP-1 agonists and SGLT2 inhibitors) targeting central mechanisms, our approach paves the way for future pharmaceutical developments. Furthermore, while the SCN is known to influence leptin directly [38] leptin receptor-resistant mice allowed us to exclude leptin-mediated mechanisms, concentrating primarily on the SCN-mediated sympathetic pathway.

In conclusion, our study shows that therapies to correct circadian misalignment could prove advantageous in treating diabetes and its related complications. Moreover, we found that correcting the central clock protects sympathetic nervous system activation and glycemic control. Collectively, these findings present a novel direction for using circadian clock genes as therapeutic targets for diabetes and related complications.

## Authors Contributions

NM and QL initiated and led the animal experiments and wrote, reviewed, and edited the manuscript. JL led the AAV experiments and edited the manuscript. QL, NM and SA obtained and analyzed data and edited the manuscript. AB edited and reviewed the manuscript. AM reviewed the manuscript. All authors have given final approval for the version to be published.

## Acknowledgment

We want to thank Dr. Charlie Dong for the helpful discussion on liver studies, Ms. Kara Orr and Lata Udari for their technical help with GTT and ITT studies, Dr. Amy S Porter, Michigan State University, for the experimental help with tyrosine hydroxylase staining, and Dr. W. Clay Smith, University of Florida, for help with AAV-Bmal design.

## Funding

This work is supported by funding support from National Eye Institute grants R01EY027779, R01EY027779-S1, and R01EY032080 to AB, a Challenge grant from Research to Prevent Blindness (RPB) to the Department of Ophthalmology.

## Conflict of Interest

AB is an *ad hoc* District Support Pharmacist at CVS Health/Aetna. The contents of this study do not reflect those of CVS Health/Aetna. NM, QL, JL and SA do not have any conflicts to declare.

## References

1. Welsh DK, Takahashi JS, Kay SA (2010) Suprachiasmatic nucleus: cell autonomy and network properties. Annu Rev Physiol 72:551–577. 10.1146/ANNUREV-PHYSIOL-021909-135919

2. Takahashi JS (2016) Transcriptional architecture of the mammalian circadian clock. Nature Reviews Genetics 2016 18:3 18(3):164–179. 10.1038/nrg.2016.150

3. Rudic RD, McNamara P, Curtis AM, et al (2004) BMAL1 and CLOCK, Two Essential Components of the Circadian Clock, Are Involved in Glucose Homeostasis. PLoS Biol 2(11):e377. 10.1371/JOURNAL.PBIO.0020377

4. Lee J, Kim MS, Li R, et al (2011) Loss of Bmal1 leads to uncoupling and impaired glucose-stimulated insulin secretion in β-cells. Islets 3(6):381. 10.4161/ISL.3.6.18157

5. Marcheva B, Ramsey KM, Buhr ED, et al (2010) Disruption of the Clock Components CLOCK and BMAL1 Leads to Hypoinsulinemia and Diabetes. Nature 466(7306):627. 10.1038/NATURE09253

6. Jouffe C, Weger BD, Martin E, et al (2022) Disruption of the circadian clock component BMAL1 elicits an endocrine adaption impacting on insulin sensitivity and liver disease. Proc Natl Acad Sci U S A 119(10):e2200083119. 10.1073/PNAS.2200083119/SUPPL_FILE/PNAS.2200083119.SD06.XLSX

7. Rakshit K, Matveyenko A V. (2021) Induction of Core Circadian Clock Transcription Factor Bmal1 Enhances β-Cell Function and Protects Against Obesity-Induced Glucose Intolerance. Diabetes 70(1):143–154. 10.2337/DB20-0192

8. Prevalence Estimates for Diabetic Retinopathy (DR) | Vision and Eye Health Surveillance System | CDC. https://www.cdc.gov/vision-health-data/prevalence-estimates/dr-prevalence.html. Accessed 18 Aug 2024

9. Lundeen EA, Burke-Conte Z, Rein DB, et al (2024) Prevalence of Diabetic Retinopathy in the US in 2021. JAMA Ophthalmol 141(8):747–754. 10.1001/JAMAOPHTHALMOL.2023.2289

10. Storch KF, Paz C, Signorovitch J, et al (2007) Intrinsic circadian clock of the mammalian retina: importance for retinal processing of visual information. Cell 130(4):730. 10.1016/J.CELL.2007.06.045

11. Mathew D, Luo Q, Bhatwadekar AD (2022) Circadian rhythm disruption results in visual dysfunction. FASEB Bioadv 4(6):364. 10.1096/FBA.2021-00125

12. Baba K, Piano I, Lyuboslavsky P, et al (2018) Removal of clock gene Bmal1 from the retina affects retinal development and accelerates cone photoreceptor degeneration during aging. Proc Natl Acad Sci U S A 115(51):13099–13104. 10.1073/PNAS.1808137115/-/DCSUPPLEMENTAL

13. Bhatwadekar AD, Beli E, Diao Y, et al (2017) Conditional Deletion of Bmal1 Accentuates Microvascular and Macrovascular Injury. Am J Pathol 187(6):1426. 10.1016/J.AJPATH.2017.02.014

14. Jidigam VK, Sawant OB, Fuller RD, et al (2022) Neuronal Bmal1 regulates retinal angiogenesis and neovascularization in mice. Communications Biology 2022 5:1 5(1):1–11. 10.1038/s42003-022-03774-2

15. Luo Q, Leley SP, Bello E, Dhami H, Mathew D, Dilip Bhatwadekar A (2022) Dapagliflozin protects neural and vascular dysfunction of the retina in diabetes. BMJ Open Diab Res Care 10:1161. 10.1136/bmjdrc-2022-002801

16. Shi SQ, Ansari TS, McGuinness OP, Wasserman DH, Johnson CH (2013) Circadian disruption leads to insulin resistance and obesity. Curr Biol 23(5):372. 10.1016/J.CUB.2013.01.048

17. Jouffe C, Weger BD, Martin E, et al (2022) Disruption of the circadian clock component BMAL1 elicits an endocrine adaption impacting on insulin sensitivity and liver disease. Proc Natl Acad Sci U S A 119(10):e2200083119. 10.1073/PNAS.2200083119/SUPPL_FILE/PNAS.2200083119.SD06.XLSX

18. Zhou B, Zhang Y, Zhang F, et al (2014) CLOCK/BMAL1 regulates circadian change of mouse hepatic insulin sensitivity by SIRT1. Hepatology 59(6):2196–2206. 10.1002/HEP.26992

19. Thackeray JT, Beanlands RS, DaSilva JN (2012) Altered sympathetic nervous system signaling in the diabetic heart: emerging targets for molecular imaging. Am J Nucl Med Mol Imaging 2(3):314

20. Christensen NJ, Brandsborg O (1974) Plasma Norepinephrine and Epinephrine in Untreated Diabetics, During Fasting and After Insulin Administration. Diabetes 23(1):1–8. 10.2337/DIAB.23.1.1

21. Krumeich LN, Cucchiara AJ, Nathanson KL, et al (2021) Correlation Between Plasma Catecholamines, Weight, and Diabetes in Pheochromocytoma and Paraganglioma. J Clin Endocrinol Metab 106(10):e4028–e4038. 10.1210/CLINEM/DGAB401

22. Rao F, Zhang L, Wessel J, et al (2007) Tyrosine hydroxylase, the rate-limiting enzyme in catecholamine biosynthesis: Discovery of common human genetic variants governing transcription, autonomic activity, and blood pressure in vivo. Circulation 116(9):993–1006. 10.1161/CIRCULATIONAHA.106.682302

23. Daubner SC, Le T, Wang S (2011) Tyrosine Hydroxylase and Regulation of Dopamine Synthesis. Arch Biochem Biophys 508(1):1. 10.1016/J.ABB.2010.12.017

24. Gao X, Wei Y, Sun H, et al (2023) Role of Bmal1 in Type 2 Diabetes Mellitus-Related Glycolipid Metabolic Disorder and Neuropsychiatric Injury: Involved in the Regulation of Synaptic Plasticity and Circadian Rhythms. Mol Neurobiol 60(8):4595–4617. 10.1007/S12035-023-03360-5/METRICS

25. Mieda M, Ono D, Hasegawa E, et al (2015) Cellular clocks in AVP neurons of the SCN are critical for interneuronal coupling regulating circadian behavior rhythm. Neuron 85(5):1103–1116. 10.1016/J.NEURON.2015.02.005

26. Vancura P, Oebel L, Spohn S, et al (2021) Evidence for a dysfunction and disease-promoting role of the circadian clock in the diabetic retina. Exp Eye Res 211:108751. 10.1016/J.EXER.2021.108751

27. Ye S, Wang Z, Ma JH, et al (2023) Diabetes Reshapes the Circadian Transcriptome Profile in Murine Retina. Invest Ophthalmol Vis Sci 64(13). 10.1167/IOVS.64.13.3

28. Ling F, Zhang C, Zhao X, Xin X, Zhao S (2023) Identification of key genes modules linking diabetic retinopathy and circadian rhythm. Front Immunol 14:1260350. 10.3389/FIMMU.2023.1260350/BIBTEX

29. Peng X, Fan R, Xie L, et al (2022) A Growing Link between Circadian Rhythms, Type 2 Diabetes Mellitus and Alzheimer’s Disease. International Journal of Molecular Sciences 2022, Vol 23, Page 504 23(1):504. 10.3390/IJMS23010504

30. Grosbellet E, Dumont S, Schuster-Klein C, et al (2016) Circadian phenotyping of obese and diabetic db/db mice. Biochimie 124:198–206. 10.1016/J.BIOCHI.2015.06.029

31. Storch KF, Paz C, Signorovitch J, et al (2007) Intrinsic circadian clock of the mammalian retina: importance for retinal processing of visual information. Cell 130(4):730. 10.1016/J.CELL.2007.06.045

32. La Fleur SE, Kalsbeek A, Wortel J, Buijs RM (2000) Polysynaptic neural pathways between the hypothalamus, including the suprachiasmatic nucleus, and the liver. Brain Res 871(1):50–56. 10.1016/S0006-8993(00)02423-9

33. Hyun U, Sohn JW (2022) Autonomic control of energy balance and glucose homeostasis. Experimental & Molecular Medicine 2022 54:4 54(4):370–376. 10.1038/s12276-021-00705-9

34. Lin EE, Scott-Solomon E, Kuruvilla R (2021) Peripheral innervation in the regulation of glucose homeostasis. Trends Neurosci 44(3):189. 10.1016/J.TINS.2020.10.015

35. Hurr C, Simonyan H, Morgan DA, Rahmouni K, Young CN (2019) Liver Sympathetic Denervation Reverses Obesity-Induced Hepatic Steatosis. J Physiol 597(17):4565. 10.1113/JP277994

36. Terazono H, Mutoh T, Yamaguchi S, et al (2003) Adrenergic regulation of clock gene expression in mouse liver. Proc Natl Acad Sci U S A 100(11):6795. 10.1073/PNAS.0936797100

37. Jouffe C, Weger BD, Martin E, et al (2022) Disruption of the circadian clock component BMAL1 elicits an endocrine adaption impacting on insulin sensitivity and liver disease. Proc Natl Acad Sci U S A 119(10):e2200083119. 10.1073/PNAS.2200083119/SUPPL_FILE/PNAS.2200083119.SD06.XLSX

38. Kalsbeek A, Palm IF, La Fleur SE, et al (2006) SCN outputs and the hypothalamic balance of life. J Biol Rhythms 21(6):458–469. 10.1177/0748730406293854

