## Supplementary material for "*BMAL1* Overexpression in Suprachiasmatic Nucleus Protects from Retinal Neurovascular Deficits in Diabetes": Supplmental Figure

**Supplementary Figures**

#### Supplementary Figure 1

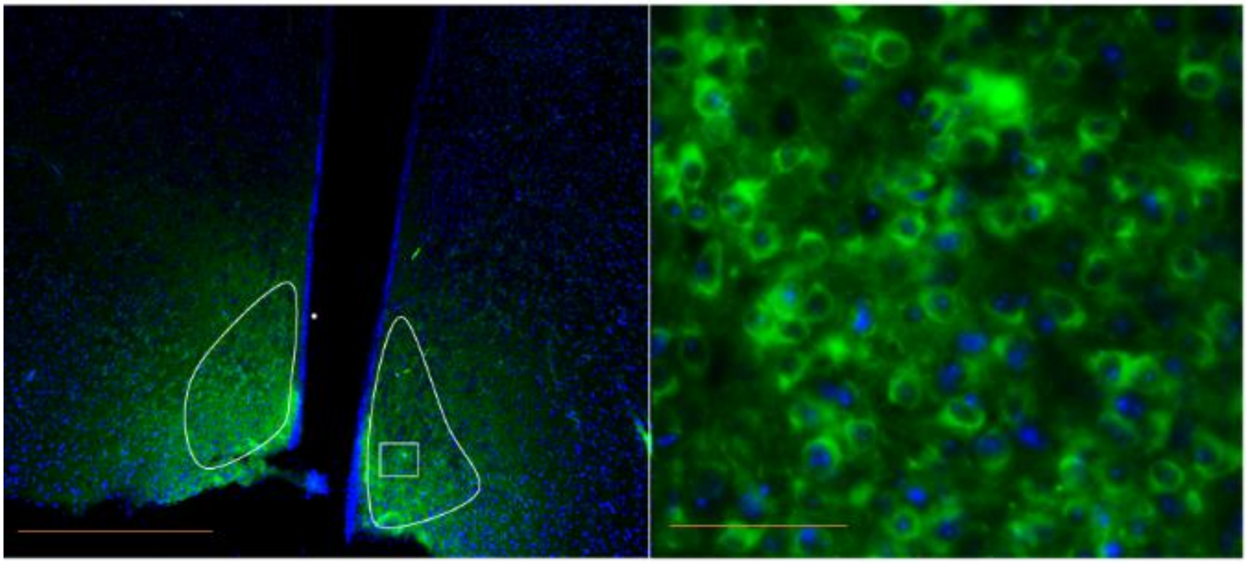

**Figure 1**: **Expression of AAV-*BMAL1* in the SCN of hypothalamus.** Mice were injected with AAV-*BMAL1* stereotaxically, four weeks post-injections, SCN region of hypothalamus were stained to locate the overexpression of *BMAL1*. Representative images showing *BMAL1* expression in the SCN. Magnification: 20X and Scale Bar - 100µM. N= 5

### Supplementary Figure 2

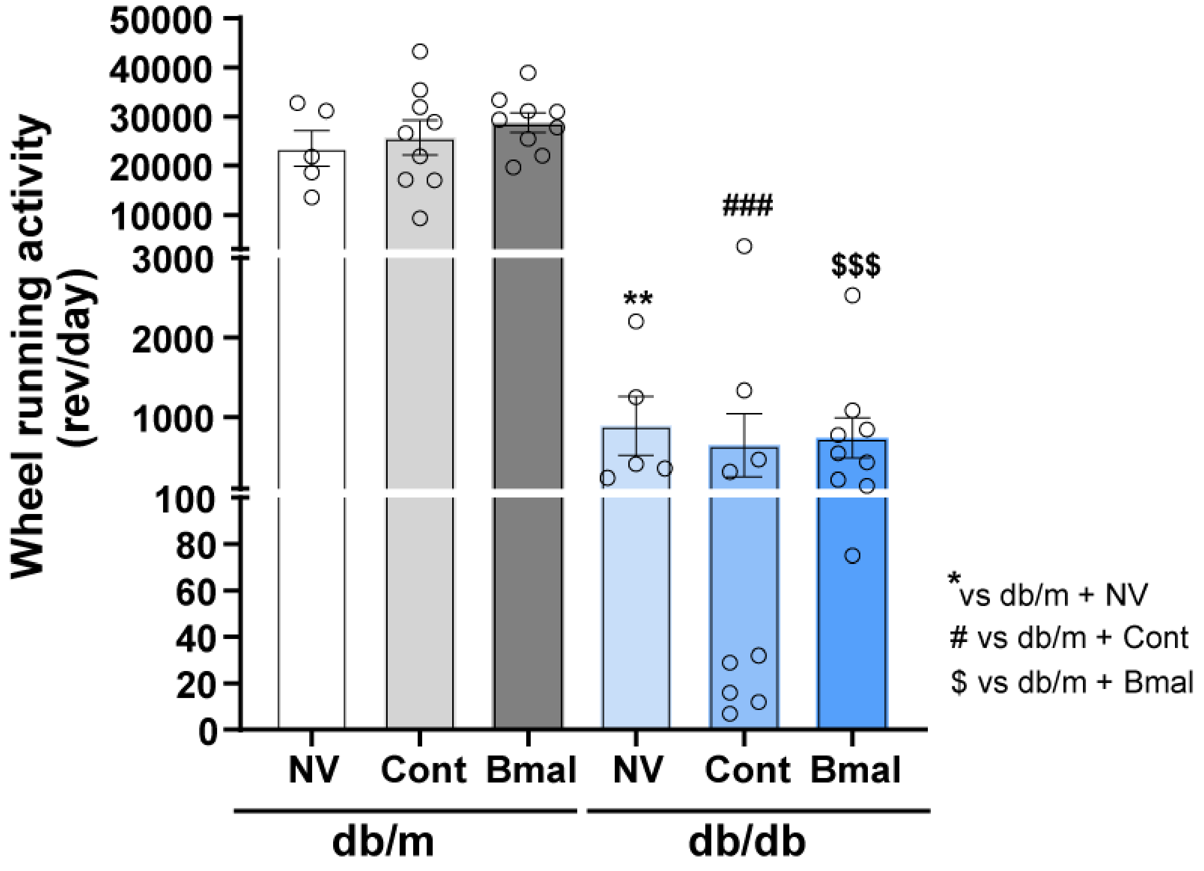


**Figure 2: Effect of *BMAL1* overexpression on wheel running activity.** Bar chart showing total wheel running activity under different conditions. N: db/m+NV-5; db/m+cont-8; db/m+*BMAL*-8; db/db+NV-4; db/db+Cont-6; db/db+*BMAL*-9. The data is presented as Mean ± SEM and analyzed using One-way ANOVA followed by Brown-Forsythe and Welch test; *$p<0.05$, **$p<0.01$, ***$p<0.001$.

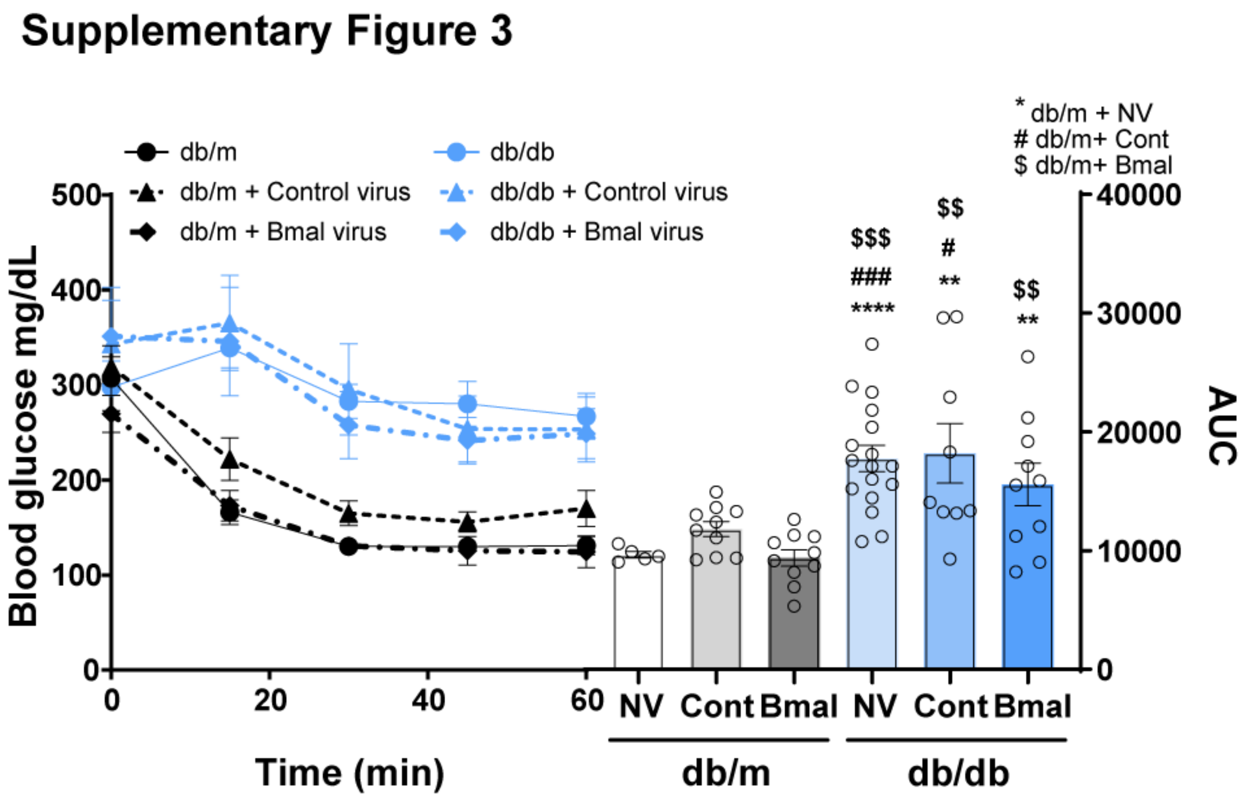


**Figure 3: Effect of *BMAL1* overexpression on insulin sensitivity in db/db mice.** The mice were fasted for 2 hours followed by an i.p. injection of insulin @ 0.5 IU/kg/bw. Blood glucose was monitored using a commercially available glucometer. ITT and its AUC in respective groups. db/m+NV-5; db/m+cont-10; db/m+*BMAL*-10; db/db+NV-12; db/db+Cont-9; db/db+*BMAL*-10.; The data is presented as Mean ± SEM and analyzed using One-way ANOVA followed by Brown-Forsythe and Welch test; * vs. db/m + NV, **p<0.01, ***p<0.001; # vs. db/m + Cont, #p<0.05, ###p<0.001; $ vs db/m + *BMAL*, $$ p<0.01, $$$ p<0.001.
